# The role of telomere shortening in carcinogenesis: A hybrid stochastic-deterministic approach

**DOI:** 10.1101/218537

**Authors:** Ignacio A. Rodriguez-Brenes, Natalia L. Komarova, Dominik Wodarz

**Author notes:** Corresponding author. (IR).

## Abstract

Genome instability is a characteristic of most cancers, contributing to the acquisition of genetic alterations that drive tumor progression. One important source of genome instability is linked to telomere dysfunction in cells with critically short telomeres that lack p53-mediated surveillance of genomic integrity. Here we research the probability that cancer emerges through an evolutionary pathway that includes a telomere-induced phase of genome instability. To implement our models we use a hybrid stochastic-deterministic approach, which allows us to perform large numbers of simulations using biologically realistic population sizes and mutation rates, circumventing the traditional limitations of fully stochastic algorithms. The hybrid methodology should be easily adaptable to a wide range of evolutionary problems. In particular, we model telomere shortening and the acquisition of two mutations: Telomerase activation and p53 inactivation. We find that the death rate of unstable cells, and the number of cell divisions that p53 mutants can sustain beyond the normal senescence setpoint determine the likelihood that the first double mutant originates in a cell with telomere-induced instability. The model has applications to an influential telomerase-null mouse model and p16 silenced human cells. We end by discussing algorithmic performance and a measure for the accuracy of the hybrid approximation.

## Introduction

Cancer is driven by a process of clonal evolution, which involves the sequential accumulation of mutations that ultimately allow for uncontrolled cell proliferation [1, 2]. Often, tumors develop different types of genome instability, which impact the tumor’s ability to evolve and progress. One important source of genome instability is telomere dysfunction [3]. While mathematical modeling has significantly advanced our understanding of tumor evolution [4], the role of telomere shortening in connection to genome instability and carcinogenesis remains poorly understood from a quantitative perspective.

A serious obstacle in modeling tumor evolution in general, is that traditional fully stochastic algorithms, such as Gillespie’s method [5], are ill-equipped to deal with population sizes that are biologically relevant to the study of tumorigenesis at the scale of cell populations. Moreover, the low mutation rates of mammalian cells require a very large number of simulations to obtain statistically meaningful results on mutant dynamics. As a consequence, too often models are constructed and analyzed with population sizes that are unrealistically small and mutation rates that are unrealistically large. This is especially problematic when trying to compare model results to emerging clinical data. Here we draw on ideas related to the development of hybrid stochastic-deterministic methods to circumvent the aforementioned limitations of fully stochastic approaches. In particular, we outline an efficient hybrid stochastic-deterministic algorithm that allows for the use of realistic population sizes and mutation rates. This algorithm should be easily adaptable to a wide range of applications in the field of evolution.

In this article, we develop a mathematical model that takes into account the effects of telomere shortening in a clonal cell population. It examines the relative likelihood and frequency of the order of acquisition of the two crucial mutations in carcinogenesis, telomerase activation and p53 inactivation, as a function of key biological parameters. We also present results on the probability that the first double mutant originates in a cell with genome instability caused by telomere dysfunction. This probability is particularly important because cells that undergo telomere-induced genome instability typically acquire a large number of genome abnormalities associated with cancer [6], which suggests that an evolutionary pathway that includes transient telomere deficiency can facilitate malignant progression [3]. To implement the model we used the hybrid stochastic-deterministic algorithm. We also discuss a measure for the accuracy of the hybrid approximation, and compare algorithmic performance to a fully stochastic implementation of the model.

## Telomeres and telomere crisis

Telomeres are repetitive sequences of DNA found at the ends of linear chromosomes. They play a protective role by hiding the chromosome ends from the DNA damage response machinery. In cells that lack telomere maintenance pathways telomere length shortens with each cell division. If cell cycle checkpoints are intact, critically short telomeres halt cell proliferation, inducing either a terminal state of arrest called cellular senescence, or apoptosis [6]. Thus, normal cells that lack telomere maintenance pathways are only capable of a limited number of divisions, a phenomenon known as Hayflick’s limit [7]. Telomerase is a ribonucleoprotein enzyme that extends telomere length. It is composed of a catalytic component that includes the protein TERT, and the RNA component TERC. Cells that express telomerase at sufficient levels offset the telomere shortening that occurs during cell division, which allows them to bypass replicative limits and divide indefinitely [3]. Since most mutations occur during cell division, replicative limits protect against cancer, by limiting the sequential accumulation of mutations and the clonal expansion of cells.

Failure of cells with critically short telomeres to undergo senescence can result in telomere crisis. During crisis continued telomere shortening leads to telomere dysfunction increasing the chance of non-homologous end joining (NHEJ) and the fusion of one dysfunctional telomere to another. Cells with fused telomeres become dicentric, which leads to breakage–fusion-bridge cycles, and high levels of genome instability and cell death [3]. Genome instability in cells undergoing crisis can give rise to chromosome gains and losses, gene amplifications and deletions, and non-reciprocal translocations amongst other types of genomic alterations. The rare cells that escapes crisis, usually through telomerase activation, typically harbor a large number of genomic abnormalities associated with cancer [6]. It has thus been suggested that the passage and emergence from crisis can be an important contributor to tumor development in some cancers [8].

In this article we use mathematical models to study the emergence and population dynamics of cells with two types of mutations: loss of p53 function and telomerase activation. Inactivation of p53 is a frequent event in tumorigenesis [9]. And in particular, inactivation of the p53 pathway is necessary to bypass telomere-induced senescence [10]. In the paper we focus on the first emergence of a double mutant and in the order of acquisition of the two mutations. We model the effects of telomere crisis by assuming an elevated death rate for unstable (in crisis) cells. The order of mutations is important, because cells that undergo crisis can acquire a number of important genomic changes, which occur during the period of genome instability caused by telomere dysfunction.

Our model has a direct application to the important TERC^−/−^ mouse model. Mouse cells have very long telomeres and express telomerase promiscuously; as a consequence telomere shortening is not a barrier to tumor progression in mice [11]. To test the function of telomerase in tissue biology a telomerase-knockout mouse model was developed, by breeding mice that do not express TERC (the RNA component of telomerase). Continuous breeding of TERC^−/−^ mice over successive generations led to the progressive shortening of telomeres [12]. A series of studies were then conducted in late generation TERC^−/−^ mice, in which a gene (Ink4a/Arf) encoding for two distinct tumor suppressor proteins was deleted. Mice null for this gene develop sarcomas and lymphomas with short latency; TERC^−/−^ mice however, had reduced tumor incidence and increased latency, demonstrating that telomere shortening and lack of telomerase expression inhibits tumorigenesis in late generation TERC^−/−^ mice [13, 14].

Critically short mouse telomeres induce senescence by activating p53; and the loss of p53 function in mice is sufficient to bypass senescence [15]. Studies of TERC^−/−^ p53^+/-^ mutant mice also revealed that the p53^+/-^ phenotype is sufficient to abrogate the normal growth arrest that occurs in response to short telomeres [16]. Neoplastic lesions in these mice had a large number of genomic aberrations consistent with telomere dysfunction and the breakage–fusion-bridge cycles that occur during crisis.

Our model also has applications to human cells that lack p16 function. In humans, stem cells, germ cells, and the vast majority of cancer cells (~ 90%) express telomerase, whereas other cell types do not [17]. The critical component of telomerase that is missing in most human cells is the catalytic subunit TERT. Unlike murine cells, human cells can trigger senescence by activating the p53 or the p16/RB pathways [10]. Although there is also evidence that suggests that p16-induced senescence is not the direct consequence of telomere shortening [18]. Regardless, cells lacking p16 function may not be uncommon in *vivo* in humans, since epigenetic silencing of the p16 gene is commonly found in histologically normal human mammary epithelial cells (HMECs) [19]. Moreover, cell culture studies of HMECs repeatedly show that following the spontaneous silencing of p16, the rare cells that are able to bypass the p53 checkpoint undergo extended proliferation and eventually enter crisis [20, 21].

## Model description

We consider four types of cells, which for notation purposes we call *X,Y,Z*, and *W*, see Figure 1A. At the base of the model we have X cells, which are telomerase negative (here noted as tmase–). Telomerase null cells correspond to TERC^−/−^ cells in the context of the mouse model previously described, or TERT negative cells in the context of human somatic cells. X cells have two functioning p53 alleles (p53^+/+^). These are proliferating cells at early possibly pre-neoplastic stages of tumor development. This characterization is consistent with the understanding that in certain tumors telomere crisis is a very early event. In breast cancer for example, telomere crisis is believed to occur during progression from usual ductal hyperplasia (UDH) to ductal carcinoma in situ (DCIS) [8]. Being telomerase negative, X cells can divide only a limited number of times. To model replicative limits we assume that each cell has a replication capacity *ρ* ≥ 0. When a cell with replication capacity *ρ* > 0 divides, it produces two daughter cells with replication capacities *ρ* – 1. Cells with replication capacity *ρ* = 0 become senescent and stop dividing (Figure 1B). The maximum replication capacity in the model is denoted by *ρ*_m_.

**Figure 1.**
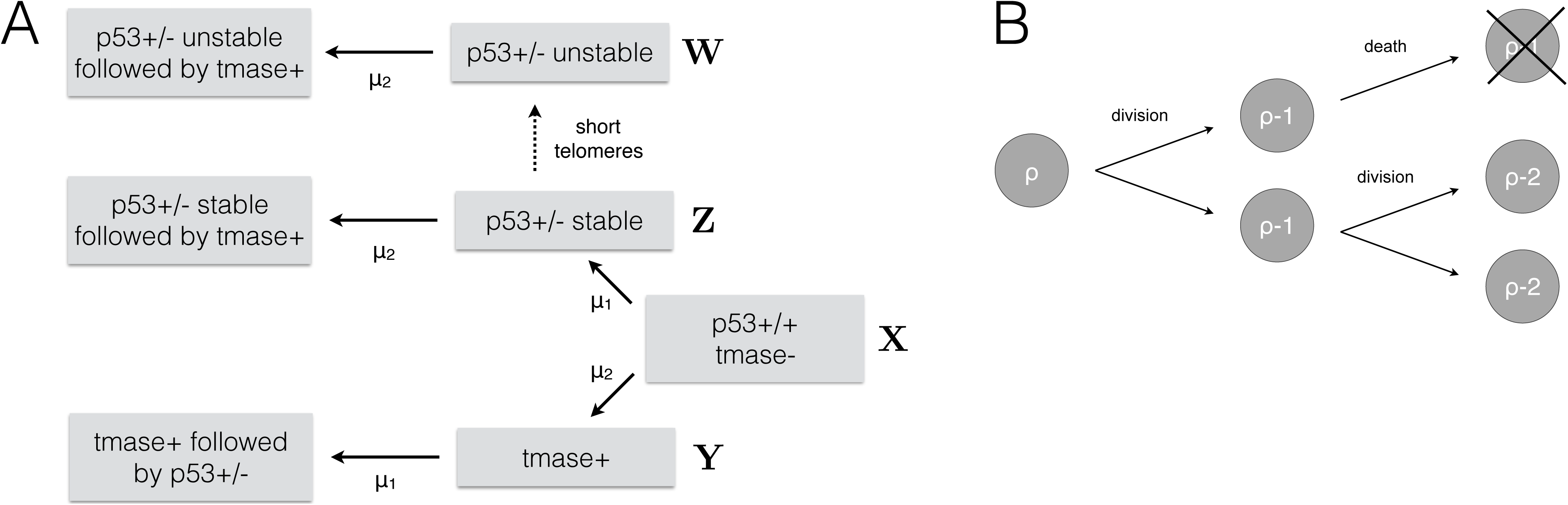
(A) Each cell has a replication capacity *ρ* ≥ 0. When a cell with replication capacity *ρ*> 0 divides, it produces two daughter cells with replication capacities *ρ* — 1. Cells with replication capacity p = 0 become senescent and stop dividing. (B) Different pathways by which cells can acquire two cancer associated mutations: Activation of telomerase (tmase+) and inactivation of one p53 allele (p53^+/-^). The mutation rate for acquiring the p53^+/-^ phenotype is set to *μ*_1_ = 10^−7^ (loss of one tumor suppressor allele). The mutation rate to activate telomerase is set to *μ*_1_ = 10^−9^ (point mutation). p53^+/-^ cells have a defective DNA damage response, which allows them to undergo extra rounds of cell division beyond the normal replication capacity p. In p53^+/-^ cells telomere length continues to decrease with each cell division, eventually leading to telomere crisis. Crisis is characterized by critically short telomeres causing chromosome breakage-fusion-bridge cycles and widespread cell death. Cells in crisis are referred in the diagram as p53^+/-^ unstable cells. Telomerase activation allows cells to escape replication limits, making them capable of dividing an unlimited number of times.

*Y* cells are telomerase positive (tmase+) and p53^+/+^. Telomerase expression allows them to escape replicative limits, making them capable of dividing an unlimited number of times. In the model, a *Y* cell arises from a point mutation in an X cell. Recently, activating point mutations in the tmase promoter have been identified in multiple cancer types [22, 23, 24, 25]. We consider mutations that occur during cell division and use the approximate point mutation rate in cancer *μ*_2_ = 10^−9^ [26].

*Z* cells are p53^+/-^ and telomerase negative. In mice, single-copy loss of p53 is sufficient to affect the cell’s ability to undergo senescence in response to critically short telomeres [16]. Direct confirmation that these same dynamics occur in humans is currently missing. However, there is strong evidence that the human p53 gene is haplo-insufficient in a wide variety of contexts [27]. Furthermore, 80% of the most common p53 mutants have been found to have the capacity to exert a dominant-negative effect over wild-type p53 [9]. Hence, in the model we assume that the p53^+/-^ phenotype allows cells to extend their replication capacity by *ρ*_e_ cell divisions beyond the point at which senescence occurs in normal cells. We call the parameter *ρ*_e_ the replication capacity extension. Early experiments, based on SV40-induced disruption of p53, suggest that the replication capacity extension is in the order of 20 PD [28], with a range of 20 to 30 PD being suggested [29]. The precise value of *ρ*_e_ however, is likely to vary *in vivo*; we thus treat it as a variable, and explore the effects of varying *ρ*_e_ on the system. In the model, *Z* cells arise from *X* cells with a rate per cell generation *μ*_1_ = 10^−7^ (a common estimate for the rate per cell division of inactivating one copy of a tumor suppressor gene [30]).

*W* cells arise from *Z* cells that keep dividing past their extended replication capacity. As a consequence their telomeres continue to shorten, up to the point where they become dysfunctional, resulting in genome instability. Cells at this stage enter crisis, a phase characterized by non-homologous end joining, breakage–fusion-bridge cycles, and widespread cell death [3]. These dynamics are considered in the model by including a separate death rate, *D*, for *W* cells.

Breast and colorectal cancer studies suggest that telomere crisis is an early event [8, 31]. In colorectal cancer, there is evidence of telomere dysfunction during the adenoma–early carcinoma transition [31]. Moreover, in a study of colorectal adenomas with average size 2 mm (range 1-3 mm) 55% of adenomas showed evidence of chromosomal instability consistent with telomere dysfunction [32]. In breast cancer, crisis is believed to occur during the UDH to DCIS transition [8], and according to a standard diagnostic criterium, ductal hyperplasias should be less than 2 mm in diameter [33]. Avascular tumors can grow up to 2-3 mm in diameter [34]. Hence, these data suggest that telomere crisis might occur during the avascular phase of tumor development. Based on these observations we limit our study to events occurring during avascular growth.

If we use a 2-3 mm diameter for avascular tumors and the volume measurements for tumor cells reported in [35], we find that the maximum cell population of an avascular tumor ranges from 3.6 × 10^6^ - 5.3 × 10^7^ cells. In the article we choose the intermediate value, *N* = 10^7^, for the maximum cell population size. To incorporate this limit in population size, we make the cell division rate dependent on cell density, controlled by the variable *f* in equation [1]. In equations [1–5], we define *K* - 10^7^/(1 — *d/r)*, where *r* and *d* are respectively the cell division and cell death rate parameters. This definition of *K* ensures that the maximum population size is equal to 10^7^, irrespective of the magnitudes of *r* and *d*; it is thus consistent with our understanding that maximum population size in avascular tumors is limited by factors such as nutrient accessibility, and not by the relative magnitudes of the cell division and cell death rates. Finally, we note that *r* and the cell death parameters, d and D, have arbitrary units of 1/time. We can then express the model in dimensionless units of time by setting *r* = 1 in the simulations and expressing the values of d and D in relation to this value of *r*.

Double mutants can be generated through a p53^+/-^ mutation in a *Y* cell (with rate **μ**_1_) or through a tmase+ mutation in a *Z* or *W* cell (with rate *μ*_2_). In the this article we are interested in the first emergence of a double mutant, for this reason when the first double mutation occurs the simulations stop. The ordinary differential equation representation of the model, including *only single* mutations (either tmase+ or p53^+/-^) is given by equations [1–5]:

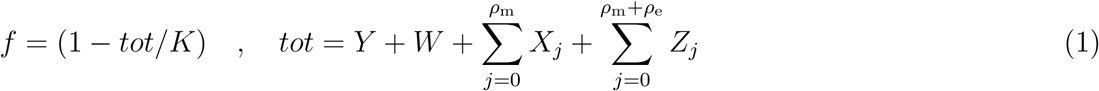

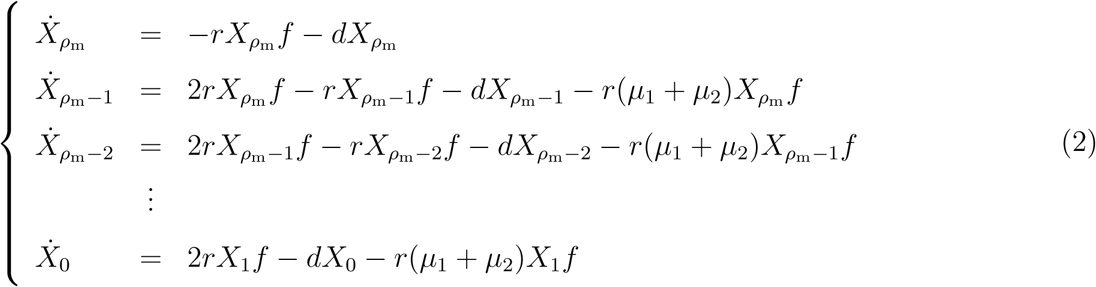

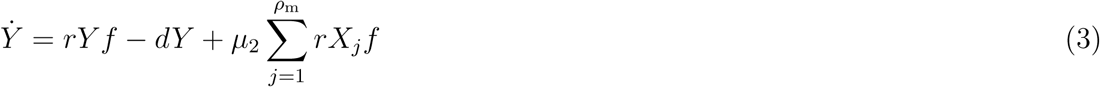

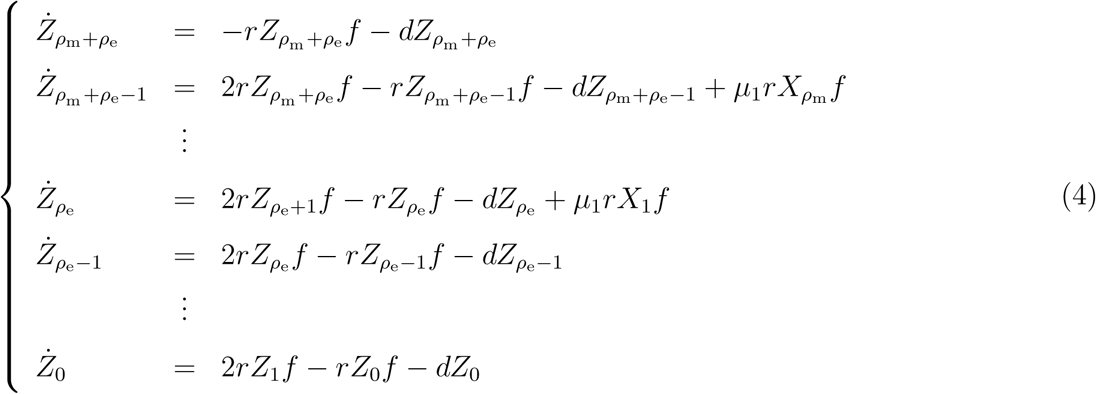

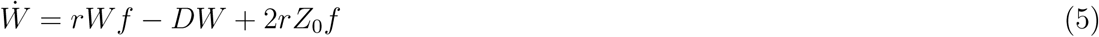

In Eqs. [1–5] we assume that both offspring of a dividing cell cannot mutate simultaneously, since the probability of such an event occurring is negligible [30].

## Hybrid method

Studying evolutionary processes computationally requires the ability to simulate the dynamics of large and small populations simultaneously. Mutations are stochastic and rare, and at least transiently, very small mutant populations can coexist with a large number of wild type individuals. In such settings, tracking the stochastic fluctuations of the small mutant populations can be essential to determine the final outcomes of the system. A problem then arises trying to simulate a multi-scale system stochastically, given that in classical fully stochastic algorithms, such as Gillespie’s method, as the population size increases the average time step decreases [5]. Recently, and especially in the field of Physical Chemistry, novel computational approaches have been developed (e.g. the Next Reaction Method and Tau-Leaping methods [36, 37]), which try to address these difficulties. There is also an important push in the development of hybrid stochastic-deterministic approaches [38, 39]. These ideas however, have not significantly penetrated the studies of population dynamics and evolution, presumably because they can rely on theoretical concepts (e.g. Langevin’s equation), which are not very common in these fields. Here, we present an application of these ideas to the field of evolution, by outlining a hybrid stochastic-deterministic algorithm for our model.

Intuitively, the implementation of the algorithm relies on two simple ideas: (i) mutations should be modeled stochastically; and (ii) if, a cell population is sufficiently large, an ODE representation can provide a good approximation of most stochastic trajectories of the population. With this idea in mind we begin with the system described in equations [1–5], which from now on we call the full system. We can write this system as a single vector equation d**V***/dt* = **F(V)**, where **V** is a vector that contains all the different cell types. Let M> 0 be a given threshold. We can classify the *X* population as small if **Σ** *X*_*i*_ <*M*, or as large otherwise, and use the same criteria to classify the other cell types *(W, Y* and *Z*). At any given time, let **V**_*l*_ and **V**_*s*_ be vectors containing the large and small cell populations. We can then define the reduced system *d***V**_*l*_/*dt* = **F**_*l*_(**V**_*l*_) derived from the full system by: (1) Retaining only the equations for the large cell populations **V**_*l*_; (2) keeping constant the contributions of the small populations **V**_*s*_; and (3) eliminating the mutation terms from the equations. If the **V**_*l*_ are sufficiently large, there will be a time interval (*t, t* + τ), where the deterministic solution of the reduced ODE will approximate the trajectories of the large populations in a stochastic implementation of the full system.

The events in the model are cell division, mutation, and death. In Gillespie’s method, every event *ν* has a given propensity *a*_*ν*_ (**V**). The time at which the next event v will occur is exponentially distributed with intensity *a*_*ν*_ (**V**). In the hybrid approach, cell division and death of large populations are modeled deterministically (using the reduced system), while cell division and death of small populations and all mutations are modeled stochastically, with propensities *a*_*ν*_ (**V**_**s**_,**V**_*l*_(*t*)) that now vary continuously with time. Hence, the next occurrence of a stochastic event v is a non-homogeneous Poisson process, with a time varying intensity *a*_*ν*_ (**V**_**s**_,**V**_*l*_(*t*)). In this case, if the system is updated up to a time *t* and *r*_*v*_ is a uniform random number in [0, 1), we can set the time for the next *ν* event as the solution, *τ*_*ν*_, to the equation [39]:

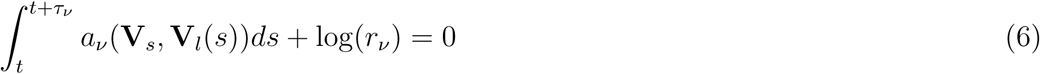

It is well known that the stochastic formulation reduces to the deterministic formulation in the thermodynamic limit [40]. However, one important practical question is how large should the threshold M be to provide a satisfactory approximation in the implementation of the hybrid algorithm. In this article, we use a numerical criterion to determine this value. First, let *G(t)* stand for the total number of cells of any of the cell types as a function of time (i.e. *G(t)* = Σ*X_i_(t),Y*(t), Σ *Z_i_(t)*, or *W*(t)). We can consider the function *E*[*G*^(*M*)^(*t*)] equal to the expected number of G type cells using the hybrid method with the threshold *M.* The *L*^2^ norm (here denoted as ||⋅||) is a measure for the distance between two functions. We can then define the normalized error *ε*(*M*_1_,*M*_2_) = || *E*[*G*^(*M*_1_^)(*t*)] − *E*[*G*^(*M*_2_)^(*t*)] ||/|| *E*[*G*^(*M*_2_)^(*t*)] ||, which provides a measure of the difference in the expected number of G cells using the two thresholds, *M*_1_ and *M*_2_, during a specific time interval *I.* To determine an acceptable threshold *M*,we define a tolerance *tol* and require that ε(*M*, 2*M)<tol*. In the result section we discuss the accuracy of the approximation for the telomere model and improvements in the computational efficiency of the hybrid algorithm compared to a fully stochastic implementation (Figure 4 and Table 1).

**TABLE I.** Execution time.

| Max Population Size | Hybrid (s) | Fully Stochastic (s) |
| --- | --- | --- |
| 10 000 | 0.10 | 0.76 |
| 100 000 | 0.11 | 6.55 |
| 1 000 000 | 0.12 | 58.47 |
| 10 000 000 | 0.22 | 503.39 |

## Results

To study the effects of replicative limits and the emergence of double mutants (p53^+/-^ and tmase+), we implement the model using a hybrid stochastic-deterministic algorithm detailed in the previous section of the paper.

Figures 2A-C plot simulations showing the three possible outcomes of the model. All simulations start with a single X type cell (tmase-, p53^+/+^) with replication capacity *ρ*_m_ = 50 (a commonly used value for human somatic cells [7]). Figure 2A depicts a simulation where a double mutation did not occur. In this panel the *X* population first rises to a value close the maximum population *(N*– 10^7^), as the replication capacity of X cells is gradually exhausted X cells stop dividing, but continue to die, which leads to their eventual extinction. During the simulation p53^+/-^ mutations take place, this allows Z cells to extend their replication capacity by *ρ*_e_ divisions. When *Z* cells exhaust their extended replication capacity, they become unstable and acquire the *W* cell phenotype, which is characterized by a high death rate *D.* Without the acquisition of a tmase+ mutation both the *Z* and *W* cell populations eventually go extinct. During this simulation tmase+ mutants do emerge (red line); however, because they do so at a time when most *X* cells have not exhausted their replication capacity they initially have no fitness advantage and in this simulation go stochastically extinct. Figure 2B depicts a simulation where a double mutant emerges from the *Y* cell population (tmase+ followed by p53^+/-^). The emergence of the double mutant is indicated by the red dot. Figure 2C plots a simulation where a double mutant emerges from the *W* cell population (p53^+/-^ unstable followed by tmase+; purple dot).

**Figure 2.**
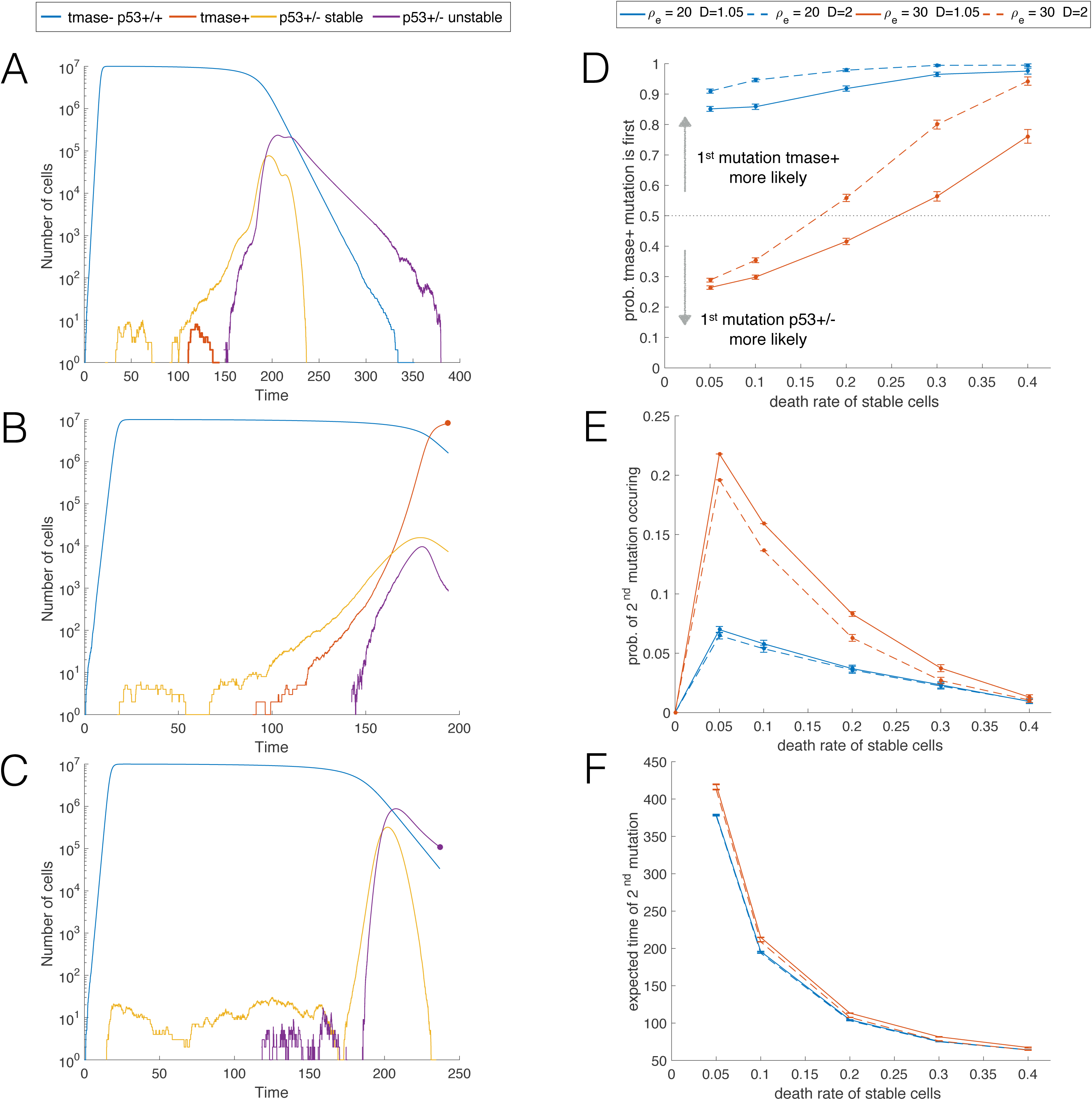
Times series of a simulation when: (A) a double mutation never occurs; (B) the first mutation emerges from *Y* cell population (tmase+ first); and (C), the first mutation emerges from the *W* cell population (p53^+/-^ first). In each panel, the first emergence of a double mutation is indicated by s solid dot. In panels A-C, *ρ*_e_ = 20, *d* = 0.1, and, *D* = 1.05. (D) Probability that the first double mutant emerges through the pathway tmase+ first followed by p53^+/-^. Error bars indicate 95% confidence intervals. Blue and red colors correspond to different values of the replication capacity extension *ρ*_e_, defined as the number of extra division that p53^+/-^ cells can undergo before entering crisis. Solid and dashed lines indicate different values D for the cell death of unstable cells (compared to a dimensionless division rate parameter *r*=1). The maximum replication capacity of of X cells (tmase-and p53^+/+^) is set to *ρ*_m_ = 50. (E) Probability of the emergence of a double mutant. (F) Expected time of the first emergence of a double mutant. Results based on 10^5^ — 10^6^ simulations per data point.

Figure 2D plots the probability that the first double mutant emerges from the *Y* cell population (tmase+ first), calculated from those simulations where a double mutation occurred. The figure includes plots for two different values of the death rate, D, of *W* cells (p53^+/-^ unstable), and two different values for the replication capacity extension, *ρ*_e_, of *Z* cells. In the model the death rate for cells in crisis (*W*) must be greater than one, otherwise cells in crisis can go on dividing indefinitely, with ever shortening telomeres and increasing levels of chromosome instability (a scenario which is not biologically feasible). For this reason, we simulated two values for the death rate of *W* cells: D = 1.05, which represents a case where the death and birth rate are nearly balanced; and D = 2 (twice the size of the birth rate parameter *r*). Figure 2D also demonstrates that the size of the replication capacity extension, *ρ*_e_, is crucial in determining the likelihood of the sequence of mutations (tmase+ followed by p53^+/-^ vs. p53^+/-^ followed by tmase+). Indeed, as shown in the simulations, a difference of only 10 cell division (*ρ*_e_ = 20 vs. *ρ*_e_ = 30) can dramatically alter the likelihood of the sequence of mutations. There is limited data for the value of *ρ*_e_, although a range of 20-30 PD has been suggested [28, 29]. The actual value of *ρ*_e_ however, is in an all likelihood cell type dependent, and influenced by multiple factors, such as the level of telomere restriction factor two (TRF2) expression [3]. Note that for *D* = 1.05 (red lines), as d increases, there is a switch from p53^+/-^ followed by tmase+ as the most likely sequence of mutations giving origin to the first double mutant, to tmase+ followed by p53^+/-^. This behavior is explained by the fact that lower values of d allow for more *Z* cell divisions, which also result in higher *W* cell populations. The higher the number of *Z* and *W* cells, the more likely that the first double mutant originates in a p53^+/-^ cell.

Figure 2E plots the probability of a double mutation occurring for different values of *ρ*_e_ and *D*. We note that the outcomes are sensitive to the value of *ρ*_e_ (red vs. blue lines). One interesting result is that when there is no cell death of stable cells *(d* = 0), the probability of a double mutation occurring is basically zero. The reason why this occurs is that tmase+ mutations are only advantageous against a background of cells that senesce and die. Otherwise *Y* cells have a neutral fitness and are thus likely to go stochastically extinct. In a setting where *X* cells die, *Y* mutants might emerge and linger on until the time when they become advantageous, but without *X* cell death, *Y* cells never gain an advantage. Here and in all figures, we performed simulations up to a maximum time *T* = 1000 (relative to a division rate parameter *r* = 1). This value of *T* was sufficient for every simulation with d> 0 to result in either complete population extinction, or the emergence of a double mutant. This would not have been the case however, if we simulated very small positive values of *d*. To understand why, we note that if the simulated time was unbounded (*T* = ∞), the probability of a second mutation occurring would be monotonically decreasing for d> 0. Indeed, as d gets smaller, the average number of *X* cell divisions increases, and thus so does the probability of a double mutant emerging. However, as d decreases, the expected time of arrival of the first double mutant goes up (Figure 2F). In fact, by the arguments in the discussion of Figure 2F, it is straightforward to see that as d goes to zero, the expected arrival time of the first double mutant goes to infinity. Hence, for any finite time interval [0, *T*], the probability of a second mutation emerging will not be monotonic for positive d, but instead will have the same basic shape as the plot in Figure 2E.

Figure 2F plots the time when a double mutation first emerges. In the simulations the mean arrival time of the first double mutant is not very sensitive to either the replication capacity extension, *ρ*_e_, or the death rate of unstable cells, D. The reason why is that mutants are not selected for until X cells start becoming senescent. As soon as the number of X cells starts declining (the time of which is unaffected by *ρ*_e_ and D), pre-existing mutant clones gain an advantage, which can lead to the arrival of the first double mutant. In the simulations as d> 0 increases, there are on average fewer cell divisions, which means that the probability of a double mutation occurring goes down (Figure 2E). Higher d values also cause *X* cells to become senescent sooner, which on average decreases the time at which mutants start to become advantageous. For this reason, even if higher values of d decrease the probability of a double mutation occurring, in those instances where a double mutation does happen, larger d values reduce the expected arrival time of the first double mutant (Figure 2F).

Figures 3A and 3B plot the probability that the first double mutant emerges from the unstable cell population (*W*), calculated from those instances where a double mutation occurred. As expected, decreasing the death rate of unstable cells increases the probability that the first double mutation originates in a *W* cell (dashed vs. solid lines). Also, increasing *ρ*_e_ by just 10 PD, from *ρ*_e_ = 20 to *ρ*_e_ = 30, significantly raises the likelihood that the first double mutant originates from an unstable cell. The dependence on d can be more nuanced. This is best exemplified by the curve corresponding to *ρ*_e_ = 30 and D = 1.05 (Figure 3B, solid line). While Figure 2D shows that the probability that the first double mutant originates in a p53^+/-^ cell goes down as d increases, it is clear from Figure 3 that the likelihood that the the first double mutant emerges from the *W* cell population can be a non-monotonic function of d. The reason behind this behavior is that smaller values of d result in more *Z* and *W* cell divisions, making the emergence of the first double mutant from a p53^+/-^ cell more likely; however, when the value of d is sufficiently small, the number of *Z* cells divisions can be large enough, so that the first double mutant can more often originate in *Z* cells directly, i.e., before p53^+/-^ cells enter crisis.

**Figure 3.**
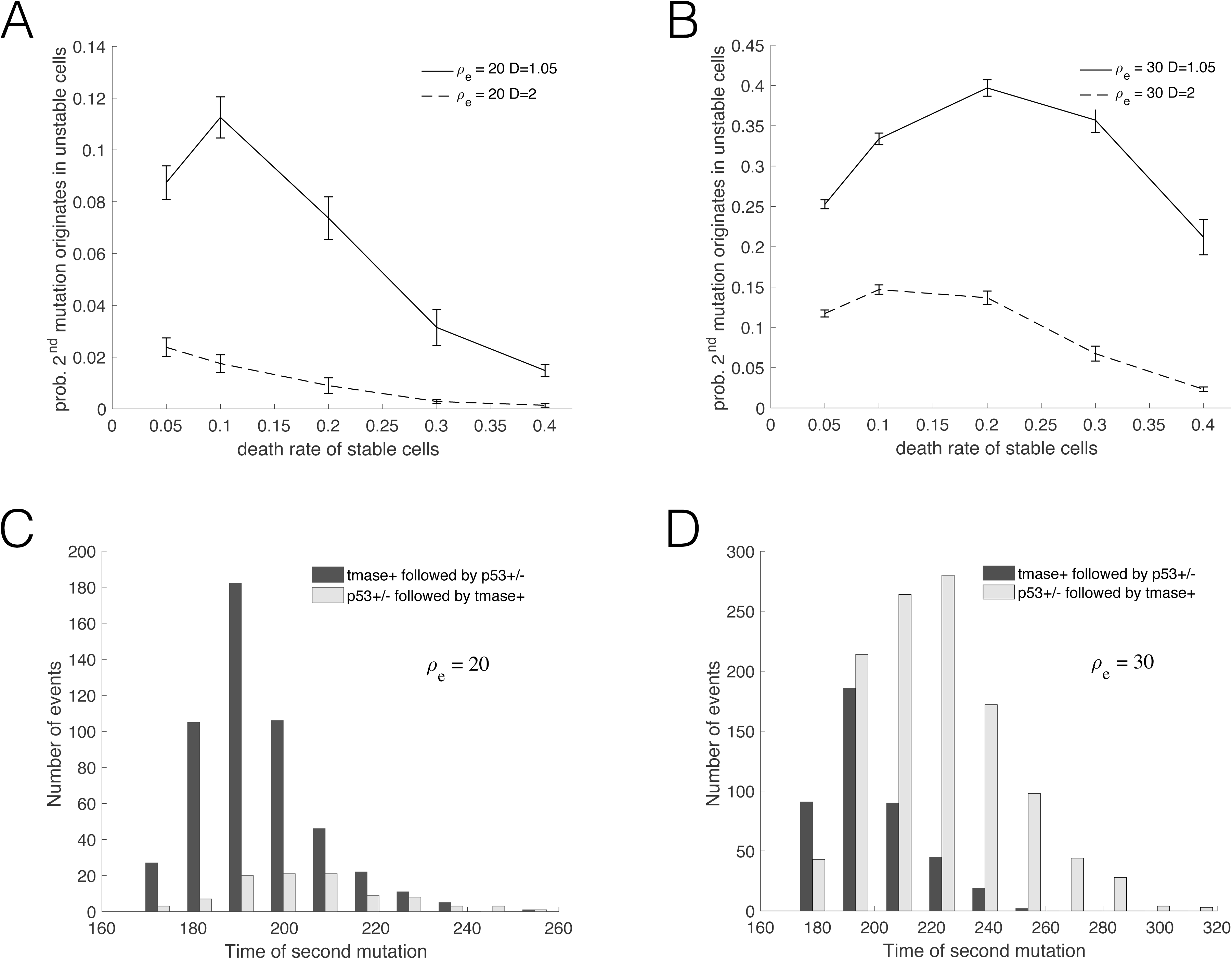
(A) and (B): Probability that the first double mutant emerges from the population of unstable (*W*) cells-conditioning over those instances where a double mutation occurred. Two different death rates of unstable cells are depicted, *D* = 1.05 (solid lines) and D = 2 (dashed lines). (C) and (D): Distribution of the arrival time of the first double mutant. The panels correspond to two different values of the replication capacity extension pe, defined as the number of extra division that p53^+/-^ cells can undergo before entering crisis. In (A) and (C) *ρ*_e_ = 20; in (B) and (D) *ρ*_e_ = 30. In all panels *ρ*_m_ = 50. In (C) and (D), d = 0.1 and *D* = 1.05.

Figures 3C and 3D present histograms depicting the distribution for the time of the first emergence of a double mutant, originating from two different sequence of events: tmase+ followed by p53^+/-^, or p53^+/-^ followed tmase+. The figure underscores the importance of the parameter *ρ*_e_ in determining the likelihood of the sequence of events. One interesting result is that, independent of the value of pe, the expected time for the emergence of the first double mutant is smaller when the second mutation originates in the *Y* cell population. In other words, the average time of emergence of the first double mutation is faster when the first mutation is tmase+.

Figure 4A plots the expected number of cells using the stochastic-deterministic thresholds *M* = 2000 (circles) and *M* = 4000 (solid lines), for simulations where double mutations did not occur-for all cell types depicted the normalized error ϵ(*M*, 2*M)*< 0.05 over the time interval I = [0, 1000]. Figure 4B plots the distribution of the times when the first double mutant emerges, using a parameter set that makes the generation of a large number of fully stochastic independent trials computationally reasonable. This figure compares the results from a fully stochastic simulation algorithm with the results from an implementation of the hybrid method. Table 1 shows the average computational run time per trial for different max population sizes using the fully stochastic and the hybrid algorithm. For a maximum population size of *N* = 10^7^ the hybrid algorithm is more than 2,200 times faster.

**Figure 4.**
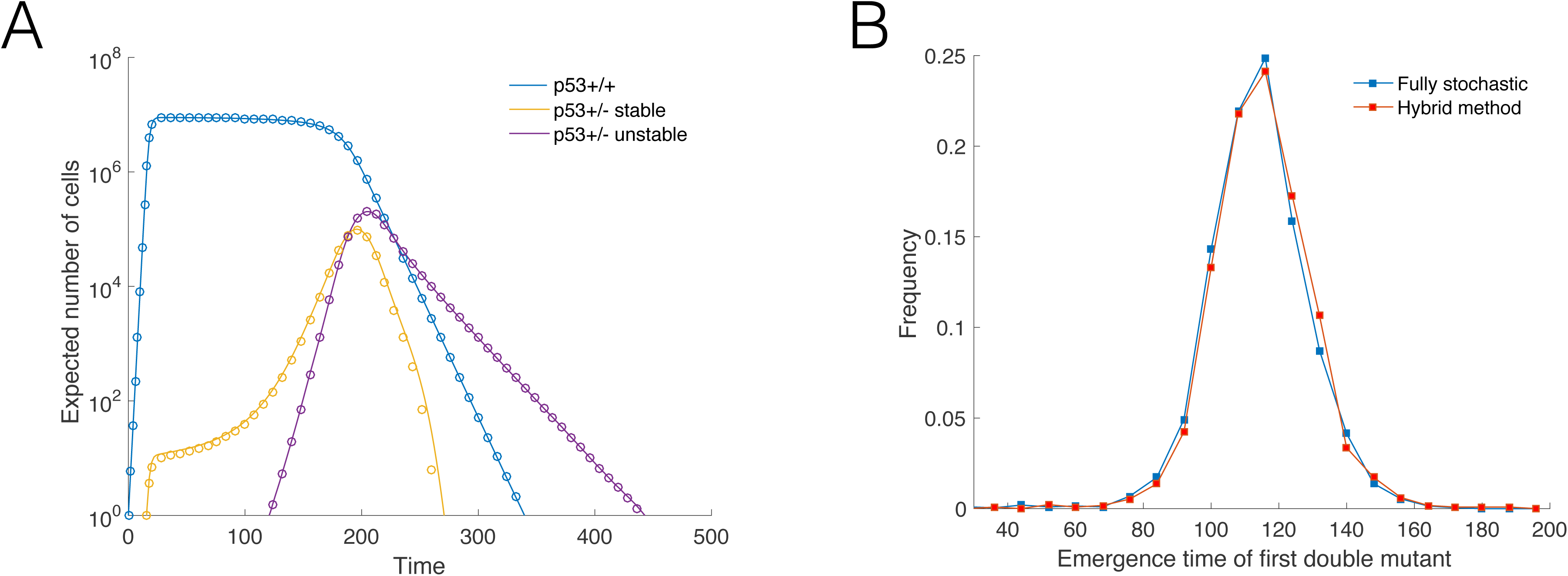
(A) Expected number of cells using two different thresholds, *M*, for the size that determines the stochastic to deterministic transition. Solid lines *M* = 2000; circles *M* = 4000. The panel corresponds to simulations where a double mutant did not emerge. Parameters: *ρ*_m_ = 50, *ρ*_e_ = 20,*d* = 0.1, and *D* = 1.05. (B) Distribution of the arrival time of the first double mutant for a parameter set that makes the generation of a large number of fully stochastic independent trials computationally reasonable. Blue: Results from fully stochastic simulations. Red: Results using the hybrid method. Parameters: *K* = 10^4^, *μ*_1_ = 10^−4^, *μ*_*2*_ = 10^−6^, *ρ*_m_ = 30, *ρ*_e_ = 15, *d* = 0.1, and *D* = 1.05.

## Discussion

Recently we presented a mathematical model with the aim of quantifying the effectiveness of replicative limits as a tumor suppressor pathway [41]. We also developed a Luria-Delbruck mutational framework to estimate the probability of escaping replicative limits through a mutation that activates telomerase [42]. These models assumed that the only constraint to cell proliferation was set by replicative limits. Here, we extend these results by studying the population dynamics in a setting where population size is also constrained by a fixed carrying capacity. We also consider the emergence of two of the most frequent events in tumorigenesis: Loss of p53 function and telomerase activation. The model has direct applications to an important telomerase negative mouse model and to p16 deficient human cells. Our work adds to growing body of literature that investigates mathematically the effects of replicative limits in cancer at the scale of cell populations (see e.g. [43, 44, 45]).

To implement our model we used a hybrid stochastic-deterministic algorithm. The algorithm simultaneously models large populations deterministically, and small populations and mutations stochastically. It provides good agreement with fully stochastic implementations of the model, and very significant improvements in terms of speed (up to several orders of magnitude faster). These improvements in performance allows us to use biologically relevant population sizes and mutation rates, circumventing some of the traditional limitations of fully stochastic methods. The development of hybrid algorithms has received considerable attention in physical chemistry applications and related fields. These ideas however, have yet to find widespread use in the field of evolution. The hybrid methodology outlined in this paper could be easily adapted to model many aspects of tumor evolution, and more broadly, it can also be applied to a wide range of evolutionary models.

In this article we examined the relative frequency of the order of acquisition of the two mutations as a function of key biological parameters. We found that for any finite time interval, the probability of a double mutation occurring is a non-monotonic function of the death rate of stable cells (d). However, if we exclude very small values of d, then increasing the death rate of stable cells decreases the probability that a double mutation occurs. Our simulations also revealed that higher death rates of stable cells increase the likelihood that the first double mutant originates in a telomerase positive cell. The probability that the first double mutant emerges from an unstable cell has a more complex dependence on d. Indeed, depending on the sizes of the replication capacity extension of p53 mutants and the death rate of unstable cells, the probability that the first double mutation originates in an unstable cell can peak at intermediate values of d. We also found that the size of the replication capacity extension of p53 mutants is crucial in determining the probability of a double mutant occurring and the likelihood of the sequence of mutations. In particular, we found that a difference of just ten population doublings in the replication capacity extension can significantly impact the behavior of the system. Interestingly, the expected arrival time of the first double mutant is only weakly dependent on the replication capacity extension and the death rate of unstable cells. Instead it is most influenced by the time at which the telomerase negative p53 wild-type cell population starts to senesce, since only then do pre-existing mutants become advantageous.

Compared to sarcomas and hematopoietic malignancies, epithelial cancers require a large number of mutations and genome rearrangements to achieve a malignant state [46]. It has thus been suggested that a mutator phenotype must take place to account for the constellation of genome abnormalities found in many malignant carcinomas. In this respect, telomere-based crisis has been identified as a key mutator mechanism driving epithelial carcinogenesis in cells that initially lack telomerase [3]. Here we presented a mathematical model that takes into account replicative limits and examines the dynamics of two mutations central to the entrance and escape from crisis. One important extension to the model will be the inclusion of mutational events, such as translocations and loss of heterozygosity (LOH), which occur at increased rates during crisis. In particular, this will require modeling the population dynamics and possible fitness differences between different types of double mutants. This analysis will be fundamental to understand quantitatively under which conditions telomere shortening shifts from being a powerful tumor suppressor pathway to a driving force behind carcinogenesis.

